# Discovery of indole-modified aptamers for highly specific recognition of protein glycoforms

**DOI:** 10.1101/2021.03.13.435263

**Authors:** Alex M. Yoshikawa, Alexandra Rangel, Trevor Feagin, Elizabeth M. Chun, Leighton Wan, Anping Li, Leonhard Moekl, Michael Eisenstein, Sharon Pitteri, H. Tom Soh

## Abstract

Glycosylation is one of the most abundant forms of post-translational modification, and can have a profound impact on a wide range of biological processes and diseases. Unfortunately, efforts to characterize such modifications in the context of basic and clinical research are severely hampered by the lack of affinity reagents that can differentiate protein glycoforms. This lack of reagents is largely due to the challenges associated with generating affinity reagents that can bind to particular glycan epitopes with robust affinity and specificity. In this work, we use a fluorescence-activated cell sorting (FACS)-based approach to generate and screen aptamers with indole-modified bases in an effort to isolate reagents that can differentiate between protein glycoforms. Using this approach, we were able to select multiple aptamers that exhibit strong selectivity for specific glycoforms of two different proteins, with the capacity to discriminate between molecules with identical tertiary structures that differ only in terms of their glycan modifications.

## Introduction

Glycosylation is one of the most abundant and diverse forms of post-translational modification. During glycosylation, covalent linkages of saccharides (*i.e.*, glycans) are attached to a target macromolecule, such as a protein or lipid. Although once thought to be of limited importance, it has become clear that glycans have a profound effect on a wide range of biological processes. For example, glycans are known to be involved in cell adhesion, molecular trafficking and clearance, receptor activation, signal transduction, and endocytosis^1^. Alterations in glycosylation have also been implicated in many human diseases^1^, and it is well known that alterations in glycosylation affect the development and progression of cancer and that such malignancy-associated glycans could be used as potential biomarkers^2^. Unfortunately, our ability to study and develop diagnostics and therapeutics based on protein glycosylation is hampered by the lack of glycan-specific affinity reagents. Anti-glycan antibodies are challenging to develop due to the inherently poor immunogenicity of carbohydrates^3^. If a glycoprotein is used as an antigen, the generated antibodies will preferentially target the protein epitopes over the glycan epitopes. Furthermore, many anti-carbohydrate antibodies exhibit poor selectivity, undermining their utility as a discriminatory reagent in the context of basic research or clinical applications^3^.

Aptamers are nucleic acid-based affinity reagents that could offer an effective alternative to antibodies for the detection of protein glycosylation. Aptamers can be selected *in vitro*, are readily chemically synthesized, and undergo facile chemical modification^3,4^. However, it has proven extremely challenging to isolate aptamers with high specificity for particular protein glycoforms, and to date there are only two examples of aptamers targeting protein glycans in the literature^5,6^. This is likely due to several factors, including the challenges of obtaining and characterizing well-defined protein glycans for use in the selection process, and the fact that aptamers generally prefer to interact with protein epitopes over glycan epitopes^3^. Despite these challenges, a DNA aptamer was recently selected that binds to a specific glycoform of prostate-specific antigen (PSA)^5^. Another group selected DNA aptamers that were chemically modified with a boronic acid moiety, which bound to the glycoprotein fibrinogen^6^. The boronic acid moieties incorporated into these DNA aptamers were shown to form key interactions with the sugar groups present on the glycoprotein. However, boronic acid groups can be problematic due to their pH-dependent binding, as well as the fact that other molecules containing *cis*-diol groups can compete with the target glycan for binding^3^. To the best of our knowledge, no other aptamer chemical modifications have been tested in the context of glycan recognition.

In this work, we demonstrate that the incorporation of the aromatic heterocyclic indole moiety into base-modified DNA aptamers can enable the highly specific recognition of protein glycan epitopes. Hydrophobic modifications have been shown to improve aptamer affinity for protein targets^7^ but have not been previously applied to glycan recognition. We identified the indole moiety as a promising candidate modification because tryptophan, an indole derivative, often forms key interactions within the binding pockets of anti-carbohydrate antibodies^8^. Using an adaptation of the multi-parameter particle display (MPPD) aptamer screening technique developed by our group^9,10^, we were able to generate aptamers bearing indole moieties that displayed exquisite glycan specificity for multiple proteins, robustly discriminating against non-target protein variants of identical structure that differ only in terms of their glycosylation.

## Results and discussion

### Strategy for selecting an N-glycan-binding aptamer

As a strategy for isolating glycan-specific base-modified aptamers, we built on our previously described MPPD technique^9,10^, which allows for the simultaneous evaluation of aptamer affinity and specificity in a single screening experiment (**Fig. 1**). MPPD aptamer selection typically begins with the conversion of a solution-phase aptamer library into monoclonal aptamer particles via an emulsion PCR process. These particles are subsequently incubated with both target and non-target molecules that have been differentially labeled with distinct fluorescent tags. These are then subjected to fluorescence-activated cell sorting (FACS) in order to identify the subpopulation of particles that generate a strong target-specific signal, but produce minimal fluorescence associated with the non-target label. We have modified this process by directly incorporating the uridine nucleotide 5-[(3-indolyl)propionamide-N-allyl]-2’-deoxyuridine-5’-triphosphate (5-indolyl-dUTP), which bears an indole moiety, as a substitute for thymidine during the emulsion PCR step. We used the commercially available KOD-XL polymerase, which has previously been shown to be capable of effectively incorporating chemically-modified nucleotides^11,12^. After the FACS screening process is complete, the collected aptamer particles are subjected to a ‘reverse-transcription’ reaction, which converts the base-modified aptamers back to natural DNA templates.

**Figure 1:**
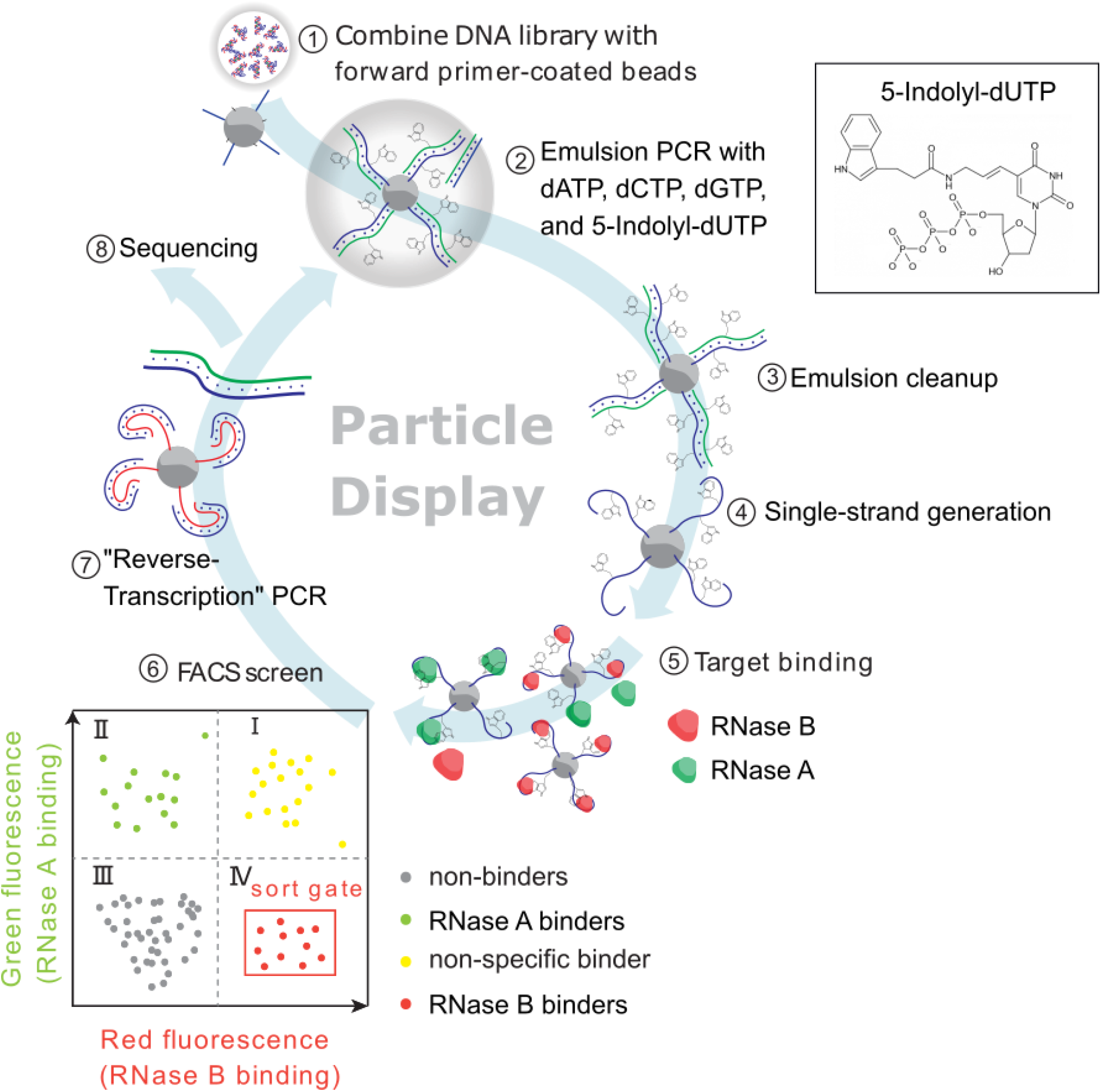
A modified multi-parameter particle display (MPPD) selection scheme for generating indole-modified aptamers. 1) DNA library molecules are hybridized to magnetic beads coated with forward primers, and 2) emulsion PCR is performed to create aptamer particles that each display many copies of a single sequence. dTTP is substituted with the modified base 5-indolyl-dUTP at this step to produce base-modified aptamers. 3) The emulsions are broken and 4) the aptamer particles are converted to single-stranded DNA via NaOH treatment. 5) The aptamer particles are incubated with RNase B (RB) and RNase A (RA), where each protein is labeled with a spectrally orthogonal fluorophore. 6) Fluorescence-activated cell sorting (FACS) separates aptamer particles that generate a strong RB-specific signal and minimal RA-specific signal (quadrant IV of the FACS plot). 7) The aptamers are subjected to a ‘reverse-transcription’ step to produce natural DNA, which is either used as the template for an additional round of screening or 8) characterized via high-throughput sequencing.

These can then be used either for a subsequent round of screening or subjected to sequencing if sufficient enrichment of target-specific aptamers has occurred.

We incorporated the indole moiety as a base-modification in the selection scheme in order to enhance the ability of the aptamer pool to form favorable contacts with glycan epitopes. It is well known that base-modified aptamers containing hydrophobic aromatic side chains can have a profound influence on the success rate of SELEX, nuclease resistance, and affinity towards protein targets (i.e. SOMAmers)^7^. X-ray crystal structures of SOMAmers bound to proteins reveal that the hydrophobic base-modification both drives the formation of unique structural motifs via intramolecular interactions, as well as participating in direct contacts with the protein^7^. Although hydrophobic base-modifications have not been previously utilized in the context of detecting protein glycosylation, we hypothesized that the indole moiety would be uniquely suited for the task due to the pivotal role that tryptophan plays in protein-carbohydrate interactions^8,13^. This is because the analysis of X-ray crystal structures of proteins non-covalently bound to carbohydrates reveal that tryptophan is highly enriched within carbohydrate-binding pockets relative to other amino acids^13^. Furthermore, it has been suggested that indole favorably binds to the electron-poor C-H bonds of carbohydrates, and that complementary electronic effects help drive protein-carbohydrate interactions^13^. Due to the established performance of SOMAmers towards protein targets, as well as the pivotal role that tryptophan plays in protein-carbohydrate interactions, we hypothesized that incorporating the indole moiety as an aptamer base-modification could greatly aid in the formation of aptamers that could specifically recognize protein glycoforms.

As an initial model for testing the advantages of selecting indole-modified aptamers for protein glycan recognition, we used RNase A (RA) and RNase B (RB). These proteins are identical in amino acid sequence, and only differ at a single N-linked glycosylation site present on RB but not on RA (**Fig. 2**). X-ray crystallography has shown that the structures of RA and RB are identical, and that the glycan present on RB does not alter the protein’s structure^14^. This is advantageous, as it allows the selection of aptamers that specifically recognize the glycan motif rather than regions of the protein that have been structurally altered as a consequence of glycosylation. Indeed, this has been suggested to be the case for the previously discovered aptamer for a glycoform of PSA^5^.

**Figure 2:**
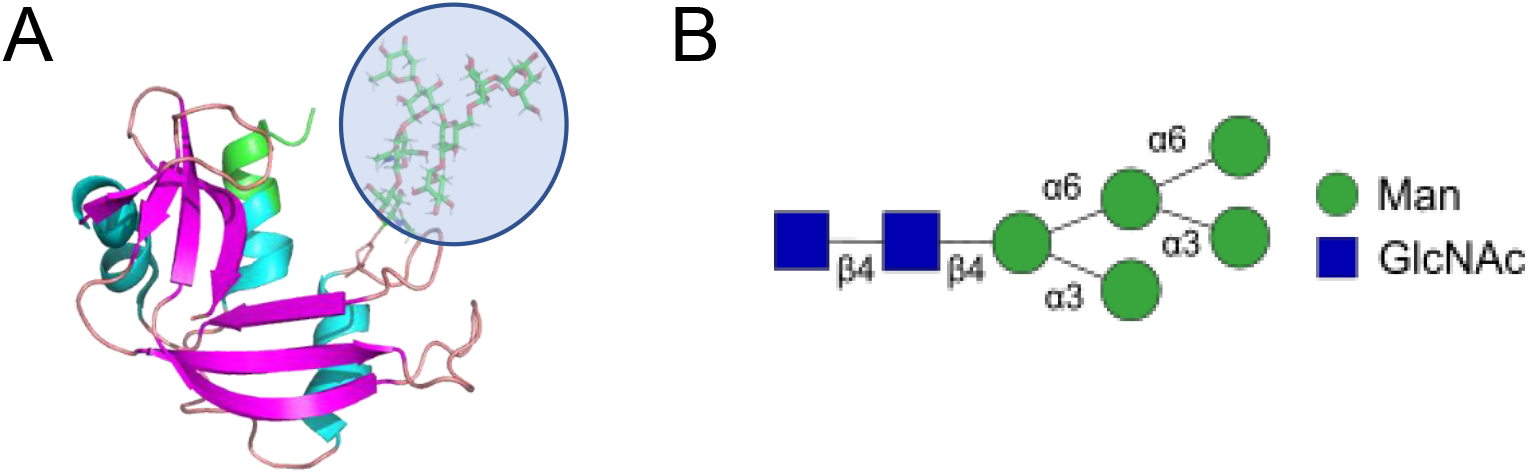
The structure of RB. **A)** The protein structure of RB, including a high-mannose N-linked glycan group highlighted in blue. The structure of RA is identical, save for the N-glycan. The figure was generated in PyMol using data from the PDB^14^ and GlyProt^15^. **B)** The N-glycan consists of mannose (Man) and N-acetylglucosamine (GlcNAc). This N-glycan is heterogeneous and can contain 1–3 additional mannose sugars via α-2 linkages^16^.

RA and RB were respectively labeled with Alexa Fluor (AF)-647 and AF-488, and then incubated with the aptamer particles for 1 hour. The samples were then washed and sorted via FACS. We then collected aptamer particles that produced increased signal in the AF-488 channel but no increase in the AF-647 channel, indicating binding to RB but not RA—and therefore, specific interaction with the glycan moiety present on RB. After a pre-enrichment step consisting of five rounds of SELEX against RB immobilized onto magnetic beads, we performed two rounds of MPPD selection. In the first round, we used 5 μM fluorescently-labeled RA and RB, which we then reduced to 500 nM in the second round to increase the stringency of the selection conditions. For both the initial library and after each round of MPPD, we performed a binding assay with the aptamer particles to assess the round-to-round enrichment of RB binders in the aptamer pool (**Fig. 3A**). For the starting library, we used 6.8 μM AF-488-labeled RB, whereas we used 3.4 μM for the MPPD-selected pools. We observed a clear increase in the fluorescence signal between the first and last round of MPPD selection, indicating increasing RB affinity as selection progressed.

**Figure 3:**
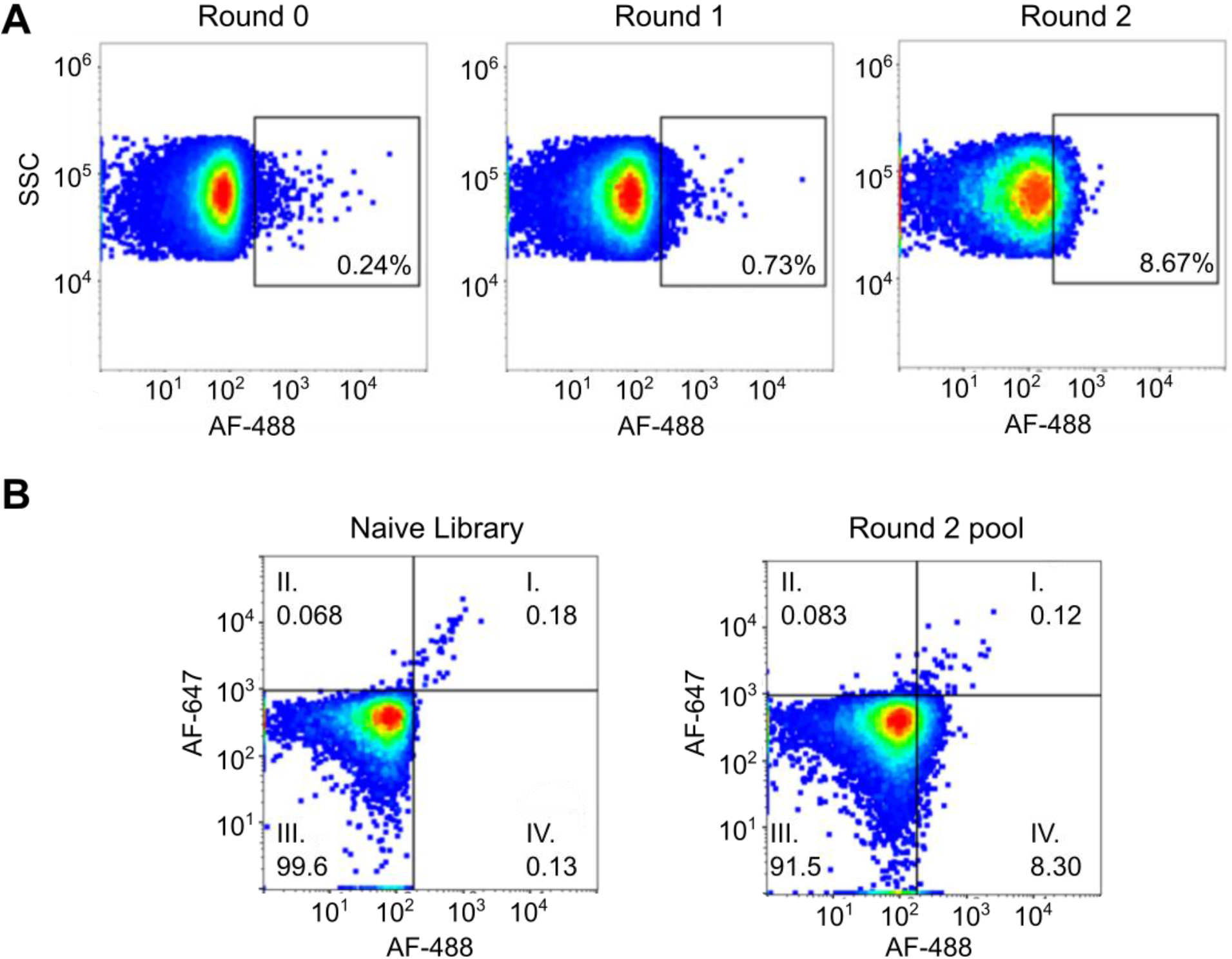
Binding assays conducted on the aptamer pools **A)** Round-to-round enrichment of the aptamer pool during particle display. Aptamer particles were incubated with fluorescently-labeled RB and analyzed via flow cytometry. The starting library (round 0) assay was conducted with 6.8 μM RB, while round 1 and 2 were conducted with 3.4 μM RB. The cytometry gates are set arbitrarily to the edge of the population and provide a consistent reference point between the plots. **B)** Two-color binding assays to simultaneously assess enrichment of aptamer pools for both RA and RB. Aptamer particles were incubated with 3.4 μM fluorescently-labeled RB and RA and analyzed via flow cytometry. Plots were generated using FlowJo.

We next performed two-channel binding assays to ensure that the enriched aptamer pool was specifically binding to the glycan epitope on RB rather than protein epitopes also present on RA. We incubated aptamer particles from either the initial library or the final aptamer pool with 3.4 μM labeled RA and RB (**Fig. 3B**). We observed a significant shift in binding to RB between the two pools, but no increase in binding to RA relative to the initial aptamer library (quadrant IV), indicating successful enrichment for glycan-specific aptamers. Likewise, the reduction in events that display strong signal in both fluorescence channels (quadrant I) likely reflects the elimination of aptamers that bind protein epitopes present on both forms of RNase.

### Characterization of RB-specific aptamer candidates

Based on the clear enrichment after two rounds of selection, we subjected the two aptamer pools to high-throughput sequencing (HTS). Aptamers were grouped into families using in-house Python code, where family members were defined as sequences having a Levenshtein distance < 4. The round 2 aptamer pool was highly converged, with the majority of the sequences falling into two aptamer families representing 45.7% and 21.5% of the total sequencing reads (**Fig. S1**). By monitoring the proportion of thymidine residues present in the aptamer sequences, we could identify enrichment of base-modified nucleotides, providing insight into their importance for glycan recognition. Although the round 2 pool did not show an overall increase in base modifications, the three most abundant families in this pool showed increased thymine content in their variable regions (30–40%). This suggests that the indole modification is an important factor enabling these aptamers to recognize the glycan motif, either directly or by forming ligand-binding structural motifs that are stabilized by hydrophobic cores.

We identified seven aptamer candidates (i-1-7), representing the most abundant sequences within the three largest aptamer families, for chemical synthesis and subsequent characterization. We utilized a bead-based fluorescence assay to measure the affinity of these aptamer candidates. Briefly, we generated aptamer particles from each aptamer candidate via bead-based PCR. These were then incubated with 20 μM fluorescently-labeled RA or RB and subjected to flow cytometry (**Figure 4A**). Both proteins were labeled with Dylight 650 to make direct comparison of binding between the two proteins clear, and to allow us to rule out the possibility that the aptamers were recognizing the AF-488 fluorophore conjugated to RB rather than the glycan motif. Each of the seven candidate aptamers showed minimal RA binding, comparable with a negative control sample that contained beads conjugated to the forward primer sequence. In contrast, all seven aptamers showed a strong and significant increase in fluorescence when challenged with RB, demonstrating that our selection procedure can isolate indole-modified DNA aptamers that selectively recognize the glycan motif of RB.

**Figure 4:**
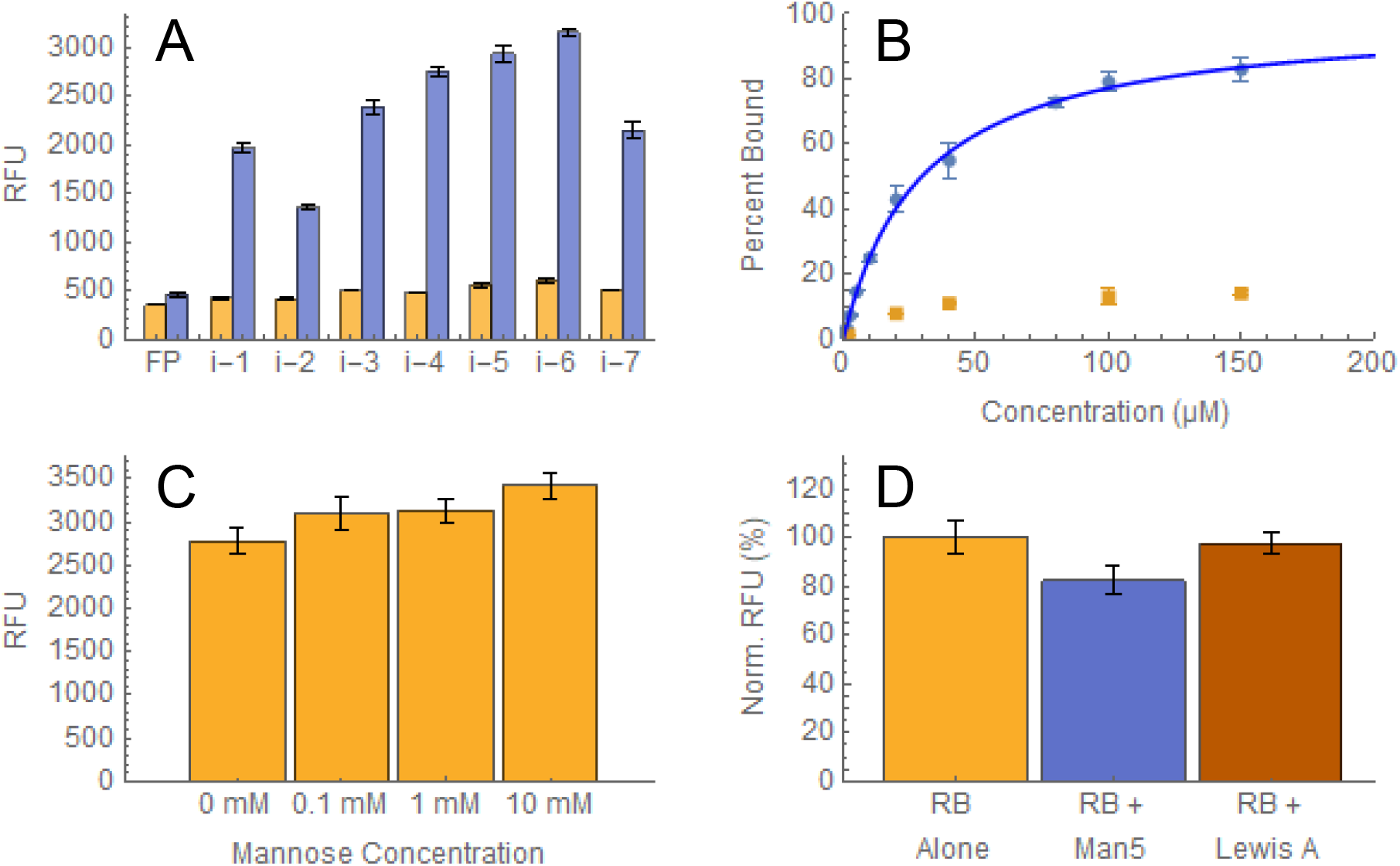
Characterization of RB aptamers. **A)** Aptamer candidates were converted to aptamer particles and incubated with 20 μM fluorescently-labeled RA (orange) or RB (blue) prior to analysis via flow cytometry. Beads containing only the forward primer (FP) were used as a negative control. **B)** Aptamer i-6 was expressed on beads and incubated with RA (orange) or RB (blue) prior to analysis via flow cytometry. The blue line represents a fitted Langmuir isotherm (*K*_*D*_ = 29.5 μM ± 2.7 μM). **C)** Competitive binding assays for i-6 aptamer particles and 20 μM fluorescently-labeled RB with mannose at concentrations ranging from 0–10 mM. **D)** Competitive binding assays conducted with i-6 aptamer particles and RB alone (orange), or with 20 μM Man5 N-glycan (blue) or 100 μM Lewis A glycan (red). The y-axis is the RFU percentage normalized to the RB-only control. All experiments were conducted in triplicate and the median RFU values were recorded. The error bars represent a single standard deviation.

Aptamer i-6 was chosen for further characterization since it displayed the strongest binding to RB of the seven aptamer candidates. We generated a full binding curve for aptamer i-6 in order to determine its *K*_*D*_. The aptamer was incubated with fluorescently-labeled RA and RB at a range of concentrations (**Figure 4B**), and fitting a Langmuir isotherm to the resulting binding curve yielded a *K*_*D*_ of 29.5 μM ± 2.7 μM. Although we observed a small increase in fluorescence at low micromolar concentrations of RA, we believe that this can be attributed to non-specific interactions as the fluorescence does not seem to further increase at even higher concentrations of RA. To demonstrate that the indole moiety was key for aptamer recognition, we synthesized aptamer i-6 with all natural nucleotides and confirmed that no binding occurred at 100 μM RB (**Fig. S2**).

We then examined the effect of enzymatically cleaving the glycan from RB on i-6 binding in order to control for unexpected differences between the two RNase variants. RB was deglycosylated with PNGase F, an enzyme that removes most N-linked glycans from glycoproteins, as confirmed by SDS-PAGE analysis. We then performed a single-concentration binding assay with RA, RB, and deglycosylated RB (**Fig. S3**). These results confirmed that removal of the glycan essentially eliminates aptamer binding, and provide additional evidence that the aptamer is specifically interacting with the glycosylated portion of RB.

Finally, we performed several competition assays to assess the glycan specificity of our newly-selected aptamer. The N-linked glycan present on RB consists primarily of mannose, and so we first assessed whether free mannose could inhibit binding between aptamer i-6 and RB due to aptamer cross-reactivity to the monosaccharide. Even at 10 mM mannose, we saw no reduction in aptamer binding to RB (**Fig. 4C**), suggesting that the aptamer does not interact exclusively with the carbohydrate moiety, but rather is recognizing a larger glycan-incorporating motif. This result highlights an advantage of using the indole moiety over the boronic acid moiety, which has a tendency to broadly and non-specifically interact with molecules containing cis-diols such as glucose, fructose, mannose, and galactose^17^. We also tested whether more complex polysaccharides could competitively inhibit aptamer binding of RB using Man5—an N-linked glycan that shares the same core structure as the high-mannose N-glycans on RB^16^. When we incubated aptamer i-6 with equimolar concentrations (20 μM) of RB and Man5, we observed a slight reduction of binding to RB (**Fig. 4D**), indicating that the aptamer can only interact weakly with this N-glycan. This could be due to a number of reasons—for example, because the glycans present on RB are heterogenous^16^, the aptamer may be interacting with protein epitopes, or due to the different conformational states of solution-phase versus protein-linked N-glycans^18^. We also observed no reduction in binding in a competition assay using the structurally dissimilar Lewis A tetrasaccharide glycan (which does not bear the common core N-linked glycan Man_3_GlcNAc_2_) at concentrations of up to 100 μM. These results collectively indicate that the aptamer preferentially binds to RB’s N-glycan motif in the context of the modified protein, and interacts minimally with free mono- or polysaccharides.

### Selection with fetuin demonstrates the generalizability of our selection platform

In order to demonstrate that this is a generalizable strategy for isolating glycosylation-specific aptamers, we conducted a selection experiment for indole-modified aptamers that specifically recognize fetuin but not asialofetuin. Fetuin is a 64 kDa glycoprotein found in the blood that is involved in multiple biological processes, such as bone remodeling and insulin resistance, and also plays a role in ischemic stroke^19^. Fetuin is heavily glycosylated, and contains sialylated N- and O-linked glycans^20^, whereas asialofetuin is created by selective removal of sialic acid from fetuin^21^ while retaining the same amino acid sequence as well as the majority of the same glycans. We anticipated that creating aptamers that distinguish fetuin from asialofetuin would be more challenging than for RNase, due to the highly similar glycan motifs shared by the two proteins.

We used the same overall selection framework as for the RB-specific aptamers, with four rounds of SELEX against bead-immobilized fetuin and counter-SELEX against asialofetuin followed by two rounds of MPPD. However, instead of directly incorporating the analog nucleotide containing the base-modification during the emulsion PCR step, we performed base modification after the PCR step using a click chemistry approach described in previous work^22^. This approach enables solid-phase synthesis of the resultant aptamer using commercially available reagents, and offers more flexibility in terms of the range of base-modifications that could be incorporated into this selection scheme. For the screening process, fetuin was labeled with Dylight 650, and asialofetuin was labeled with Dylight 532. We used 100 nM labeled fetuin in both rounds of MPPD, but only introduced 200 nM labeled asialofetuin in the second round. The resulting aptamer pool was then sequenced on an Illumina MiSeq.

We selected seven aptamer candidates and screened them via flow cytometry with 1 μM fluorescently-labeled fetuin or asialofetuin (**Fig. 5A**). All seven aptamers exhibited increased fetuin binding compared to a FP-only negative control, and minimal binding to asialofetuin. We selected aptamer f-4 for further characterization since it displayed the greatest increase in fluorescence when challenged with fetuin. After incubating aptamer f-4 with fetuin and asialofetuin at a range of concentrations, we generated a binding curve and fitted the data to a Langmuir binding isotherm (**Fig. 5B**). Based on these data, we measured a *K*_*D*_ of 6.2 μM ± 0.2 μM for fetuin, with no measurable binding observed for asialofetuin, confirming the excellent specificity that can be achieved for protein glycoforms with our selection process.

**Figure 5:**
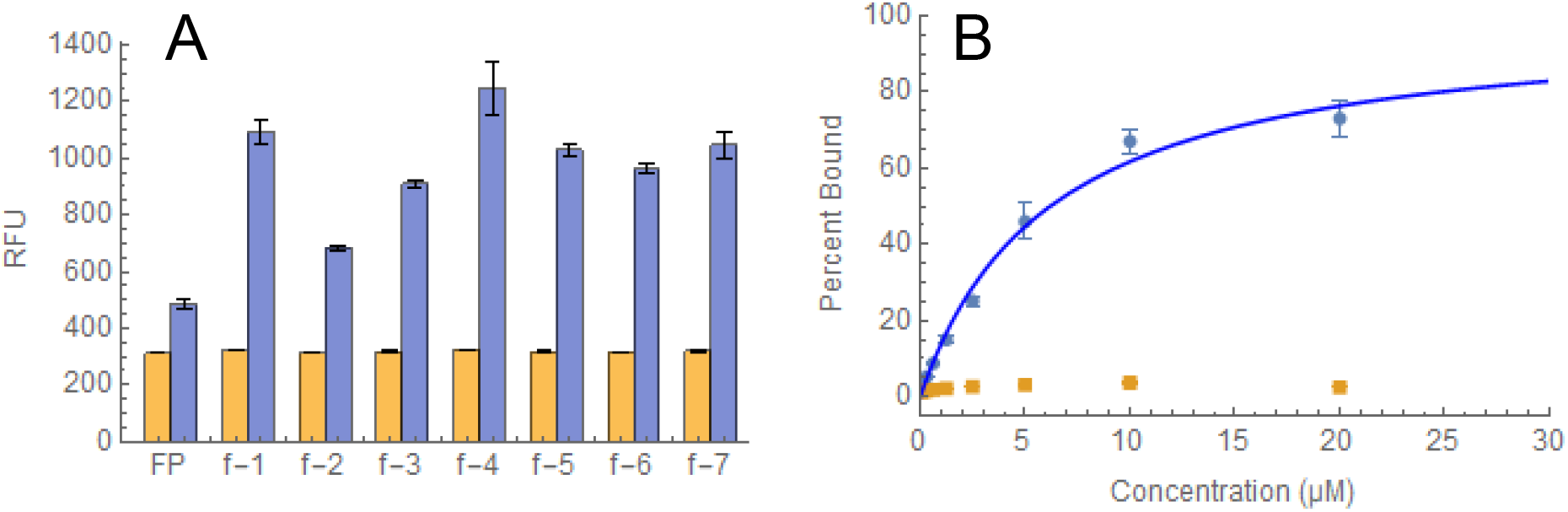
Characterization of fetuin aptamers. **A)** Aptamer particles displaying seven aptamer candidates were incubated with 1 μM fluorescently-labeled fetuin or asialofetuin prior to analysis via flow cytometry. Beads containing only FP were used as a negative control. **B)** Beads coated with aptamer f-4 were incubated with either fetuin (blue) or asialofetuin (orange) prior to analysis via flow cytometry. The blue line represents the results of a fitting a Langmuir binding isotherm to the fetuin binding data (*K*_*D*_ = 6.2 μM ± 0.2 μM). Each experiment was conducted in triplicate and the error bars represent the standard deviation.

## Conclusion

In this work, we describe a workflow for the generation of indole-modified aptamers that can recognize protein glycoforms with high specificity. We have demonstrated the generalizability of the process by performing selections against two different glycoproteins, and show that the resulting aptamers do not exhibit meaningful affinity even for protein variants of identical structure that differ by just a single glycan. Notably, our RB-specific aptamer also retains strong target affinity even in the presence of free N-glycans, indicating that this reagent only recognizes the carbohydrate group in the milieu of the fully folded protein.

Aptamers that bind strongly and selectively to specific protein glycoforms could prove extremely useful as both research and diagnostic tools. This work strongly indicates that the indole moiety offers an advantageous aptamer base-modification for glycan recognition, with advantages relative to the previously-characterized boronic acid functional group in terms of specificity. Although it is outside of the scope of this study, future work could investigate the mechanism by which indole-modified aptamers interact with carbohydrates, and if the mechanisms are similar to those involved in protein-carbohydrate interactions. This same screening approach could be used in the future to explore additional aptamer base-modifications that could aid in the generation of glycoform-specific aptamers. In particular, it may be valuable to examine alternative hydrophobic base-modifications that have been shown to aid in aptamer-protein recognition, such as benzyl-dU and napthyl-dU^23^, as well as base-modifications that are similar to other non-polar protein-residues that have been shown to stabilize protein-carbohydrate complexes (e.g. phenylalanine, alanine, methionine, leucine, proline)^7^. We selected the glycans targeted in this study because they are well characterized, which was advantageous for the creation and validation of the aptamer generation pipeline. However, we see no reason based on the results obtained here that the same workflow could not be applied to glycoprotein targets with direct clinical or diagnostic applications—for example, producing aptamers that can discriminate disease-related from healthy glycoforms. The generation of indole-modified aptamers should offer a valuable starting point towards this goal, and it is our hope that future research will reveal additional base-modifications that further expand the scope and utility of the glycobiology toolbox.

