## Supplemental information for "Discovery of indole-modified aptamers for highly specific recognition of protein glycoforms"

**SI Materials:**

N-(3-dimethylaminopropyl)-N’-ethylcarbodiimide hydrochloride (EDC) was purchased from Sigma Aldrich (#161462). N-Hydroxysuccinimide (NHS) was purchased from Sigma Aldrich (#130672). The methyl-PEG12-amino was order from Thermo Fisher Scientific (#26114). RNase A and RNase B were purchased from Sigma-Aldrich (#R6513 and #R1153). Fetuin (#F3004) and asialofetuin (#A4781) were purchased from Sigma-Aldrich. The nucleotide analog 5-indolyl-AA-dUTP was ordered from TriLink Biotechnologies (#N-2065) and the KOD XL polymerase was ordered from Thermo Fisher Scientific (#71-087-4). The C8-alkyne-dUTP was purchased from Jena Bioscience (#CLK-T05-S). Tryptophan azide was purchased from ChemPeP (#182033). Remove-iT PNGase F and (#PO706S) and chitin magnetic beads (#E8036S) were obtained from New England Biolabs (NEB). The N-glycan used in the competition assay was obtained from ProZyme (Glyko Oligomannose 5 (Man5), #GKM-002500). The Lewis A tetrasaccharide was obtained from Biosynth Carbosynth (#OL09843). The single-stranded DNA (ssDNA) library, primers, aptamer template sequences, and sequencing adaptor primers were ordered from Integrated DNA Technologies (IDT). The library for the RNase B selection was 90 nt long, consisting of a 50 nt variable region flanked by two 20-nt primer sites (5’-AGCAGCACAGAGGTCAGATG-N50-CCTATGCGTGCTACCGTGAA-3’). For the RNase library the FP sequence was 5’-AGCAGCACAGAGGTCAGATG-3’ and the RP was 5’- TTCACGGTAGCACGCATAGG-3’. The RNase sequencing adaptors were 5'-TCGTCGGCAGCGTCAGATGTGTATAAGAGACAGNNNNAGCAGCACAGAGGTCAGATG-3’ for the FP adaptor and 5'-GTCTCGTGGGCTCGGAGATGTGTATAAGAGACAGNNNN TTCACGGTAGCACGCATAGG-3’ for the RP adaptor. The library for the fetuin selection was 5’- CCAGCGAGCCAGCGAC-(VNVV)_10_-CACGCAGGACGGCACAG-3’ where V was a mixed base code for A, C, or G. The fetuin FP sequence was 5’-CCAGCGAGCCAGCGAC-3’ and the RP sequence was 5’-CTGTGCCGTCCTGCGTG-3’. The fetuin sequencing adaptors were 5’-TCGTCGGCAGCGTCAGATGTGTATAAGAGACAGHNNNNAGCAGCACAG AGGTCAGATG-3’ for the FP adaptor and 5’-GTCTCGTGGGCTCGGAGATGTGTATAAG AGACAGNNNNCTGTGCCGTCCTGCGTG-3’ for the RP adaptor. For each of the libraries 5’-amino-Spacer18 modified FP strands, 5’ fluorescein-labeled FP complement strands, 5’-biotin-RP strands, and Alexa Fluor 647 RP strands were ordered from IDT and HPLC purified.

**SI Methods:**

MPPD PCR protocol:

Unless stated otherwise, the following PCR protocol was used for all experiments: 95 °C for 5 minutes, (94 °C for 15 seconds, 54 °C for 30 seconds, 72 °C for 1 minute) x cycles, 72 °C for 5 minutes, 4 °C hold.

Conjugation of proteins to magnetic beads:

Proteins were immobilized onto MyOne carboxylic acid magnetic beads (Thermo Fisher Scientific) according to the manufacturer’s protocol for two-step coating using EDC/NHS. RA and RB were coupled to the magnetic beads using an identical two-step NHS/EDC procedure. 300 μL of magnetic beads were washed four times with 0.1% Tween-20 in PBS, and then washed twice with 300 μL of MES (pH 6), with ten minutes of incubation at room temperature (RT) on a rotator for each wash. The beads were then resuspended in 50 μL of 50 mg/mL NHS and 50 μL of 50 mg/mL EDC solution and incubated at RT on a rotator for 30 minutes. The magnetic beads were then washed twice with 300 μL cold 25 mM MES buffer (100 mM 2-(N-morpholino)ethanesulfonic acid, pH 6). The beads were then resuspended in 100 μL of RA or RB solution (2 mg/mL in PBS) and 67 μL of MES. For fetuin and asialo fetuin, beads were resuspended in 40 μL of each protein solution (10 mg/mL in PBS) 50 μL 100 mM MES, and H_2_O to a final volume of 200 μL. Samples were vortexed and incubated for 1 hour at RT on a rotator. The beads were then incubated with 500 μL of bead wash buffer for 15 minutes, washed twice with 500 μL of wash buffer, washed once with 500 μL of 0.1% Tween-20 in PBS, and then resuspended in 300 μL of 0.1% Tween-20 in PBS and stored at 4 °C.

Pre-enrichment by SELEX for RB:

Several rounds of SELEX were conducted to enrich the library and reduce the sequence space, because FACS can only screen ~10^8^ particles in a reasonable time-frame. In the first round, natural DNA was used and no counter-selection was performed. In this round, 50 μL of RB beads were washed twice with 500 μL of selection buffer (40 mM Tris-HCl, 240 mM NaCl, 10 mM KCl, 2 mM MgCl_2_, 2 mM CaCl_2_, and 0.01% Tween-20 in nuclease-free water) and incubated for 15 minutes during the second wash. The beads were then resuspended in 200 μL of selection buffer with 2.5 μM naïve library for 45 minutes. The supernatant was removed, and 100 μL of selection buffer was added. The sample was heated at 95 °C for 5 minutes to elute the bound aptamers, and then the supernatant containing the aptamer pool was collected. The natural DNA library was then converted to the base-modified DNA library in a large-scale 5 mL PCR reaction (1X KOD XL buffer, 0.1 mM dATP, 0.1 mM dGTP, 0.1 mM dCTP, 0.1 mM 5-indolyl-AA-dUTP, 250 nM 5’ biotinylated RP, 250 nM FP, 1 nM template DNA, and 250 U of KOD XL polymerase, brought up to volume with water). Seven cycles of PCR were performed, and the dsDNA was cleaned up using a Qiagen QIAquick PCR purification kit (#28104). 500 μL of Dynabeads MyOne Streptavidin C1 beads (Thermo Fisher Scientific #65002) were washed twice with 500 μL of bead wash buffer and resuspended in 1 mL of bead wash buffer (150 mM NaCl, 9.7 mM Tris, 9.7 uM EDTA, 0.1% Tween-20 in nuclease-free water, pH 7.5). The base-modified DNA was then added and incubated at RT on a rotator for at least 20 minutes. The beads were washed twice with 1 mL of wash buffer, once with 500 μL of water, and then resuspended in 100 μL of water. The sample was then heated to 95 °C for 5 minutes, and the supernatant containing the base-modified aptamer library was collected and quantified using a Nanodrop spectrophotometer. Four additional rounds of enrichment were conducted using the base-modified aptamer library with the same process described above, except 10 μL of RB beads were used in each round and the PCR reaction was scaled down to 1 mL. In the final round, counter-selection was introduced using 5 μL of RA beads.

Pre-enrichment by SELEX for fetuin:

Four rounds of positive and negative SELEX were performed with bead-immobilized fetuin and asialofetuin. All bead washing steps were performed using a Dynamag-2 magnetic separation stand. 1 nmol fetuin library was folded in 200 μL of fetuin selection buffer (100 mM NaCl, 2 mM MgCl_2_, 5 mM KCl, 1 mM CaCl_2_, 0.02% Tween 20, 20 mM Tris-HCl, pH 7.5) by heating to 95 °C for 5 min, cooling at 4 °C for 10 min, and then incubating at 25 °C for 10 min. 4 nmol (20 μl) of fetuin-conjugated beads were washed twice with 200 μl fetuin selection buffer and resuspended in the 200 μl of folded library and incubated at RT with rotation for 1 hr. Beads were washed twice with 100 μl selection buffer and eluted into 100 μl water by heating to 95 °C for 5 min twice. Recovered DNA was purified with a Qiagen MiniElute cleanup kit and eluted in 10 μl water. We performed PCR under the following reaction conditions: 10 μl 2X GoTaq PCR mix, 200 nM Fet FP, 200 nM biotinylated Fet RP, 1.5 μl recovered library, and H_2_O up to a final reaction volume of 50 μl using the following cycling conditions: 95 °C for 3 min, followed by X cycles of 96 °C for 15 s, 57 °C for 30 s, 72 °C for 30 s and finally 72 °C for 2 min. To determine the correct number of cycles for amplification, a pilot PCR was run. 5 μl of the reaction was removed every 2 cycles (16 cycles total), and then run on a 10% TBE gel at 200 V for 40 min. The cycle that yielded a product of the correct length without forming undesired products was chosen for the final scaled-up PCR reaction. To generate ssDNA, biotinylated dsDNA was immobilized onto 100 μl SA C1 Dynabeads according to the manufacturer’s protocol in 1X Binding and Washing buffer (5 mM Tris-HCl (pH 7.5), 0.5 mM EDTA, 1 M NaCl) in a 1.5 mL Eppendorf tube. Beads were incubated with 100 μl freshly-prepared 0.5 M NaOH for 10 minutes at RT. The tube was placed on a DynaMag-2 magnetic rack (Life Technologies) for 2 minutes, and the supernatant was collected. Beads were washed once more with 50 μl 0.1 M NaOH. DNA was recovered from NaOH by adjusting the pH with 25 μl 3M NaOAc, then purified with a Qiagen MiniElute cleanup kit and eluted in 20 μl water. Rounds 2-4 incorporated negative SELEX after ssDNA generation and prior to positive SELEX. The entire ssDNA library was folded as previously described, and incubated with a 10-fold molar excess of asialofetuin beads for 1 hr at RT with rotation. Beads were washed twice with 100 μl selection buffer, and the supernatant was collected and incubated with a 10-fold molar excess of fetuin beads and incubated at RT for 1 hr with rotation. DNA was recovered and amplified as described above.

Forward primer (FP) bead conjugation protocol:

500 μL of Dynabeads MyOne Carboxylic Acid magnetic beads (ThermoFisher Scientific) were washed 5 times with 500 uL of water on a magnetic rack. The beads were then resuspended in 150 μL of 0.2 mM 5’ amino-modified FP, 200 mM NaCl, 1 mM imidazole chloride and 250 mM EDC. The mixture was mixed well and sonicated prior to incubation at RT overnight on a rotator. This results in the covalent coupling of 5’-amino-Spacer18-FP to the carboxylic acid groups on the magnetic bead surface. Next, we conjugated PEG12 to the unreacted free carboxyls on the magnetic beads through a two-step NHS/EDC reaction to reduce non-specific interaction with the target proteins. The beads were washed three times with 500 μL of 100 mM MES buffer (pH 4.7). During the last wash step, the beads were incubated for 10 minutes at RT on a rotator. Immediately before use, an 80 mg/mL solution of EDC and a 25 mg/mL solution of NHS were prepared in cold 100 mM MES buffer. The FP beads were then resuspended in equal volumes of NHS and EDC solutions to a final volume of 150 μL. The beads were mixed well and incubated at RT on a rotator for 30 minutes. The beads were washed twice with 500 μL of cold PBS. The activated beads were then resuspended in 150 μL of 20 mM amino-PEG in PBS, mixed well, and incubated for at least 30 minutes at RT on a rotator. The beads were then washed three times for 15 minutes with 500 μL of wash buffer in order to quench any amine-reactive NHS esters. Finally, the beads were resuspended in 500 μL of wash buffer and stored at 4 °C.

In order to make sure FP was successfully conjugated to the beads, 1 uL of FP beads was added to 100 μL of 100 nM fluorescein-labeled FP complement and incubated for 10 minutes at RT on a rotator. The beads were then washed once with 500 μL of wash buffer, resuspended in 200 μL of wash buffer, and run on a benchtop flow cytometer (BD Accuri C6 Plus).

Emulsion PCR protocol:

The emulsion PCR process involves the creation of an oil phase and an aqueous phase. The oil phase consists of 4.5% Span-80, 0.4% Tween 80, and 0.05% Triton X-100 in mineral oil (all purchased from Sigma-Aldrich), stored at RT in the dark. The aqueous phase consists of 1X KOD XL buffer, 0.5 U of KOD XL polymerase, 0.2 mM dATP, 0.2 mM dCTP, 0.2 mM dGTP, 0.2 mM of the base-modified nucleotide, 10 nM FP, 1 μM RP, 2 pM dsDNA aptamer library, and ~3x10^8^ FP-coated magnetic beads (12 μL of FP-bead suspension) in a total volume of 1 mL of water. To create the water-in-oil emulsions, 7 mL of the oil phase were added to a DT-20 tube (IKA) and 1 mL of the acqeous phase was added dropwise over ~30 seconds while the mixture was stirred at 600 rpm in an Ultra-Turrax device (IKA). The mixture was then stirred on the device for another 5.5 min. The emulsion was then hand pipetted into ~80 wells of a 96-well PCR plate (100 μL per well). The plate was then run on a PCR machine for 40 cycles.

Emulsion cleanup:

After PCR the emulsions were transferred to a 50 mL Falcon tube. 125 μL of 2-butanol (Thermo Fisher Scientific) was added to each well that had contained emulsion, and the butanol was then transferred to the same 50 mL tube. The tube was vortexed for 30 seconds rigorously and then centrifuged at 3000 x g for 6 minutes. A magnet was used to retain the pellet of aptamer particles while removing the supernatant. 1.2 mL of breaking buffer (100 mM NaCl, 1% Triton X-100, 10 mM Tris-HCl, pH 7.5, and 1 mM EDTA) was added to the particles, and the mixture was transferred to a new 1.5 mL tube. The 1.5 mL tube was vortexed and centrifuged at 21,000 x gfor 1 minute. Using a magnetic rack, the supernatant was removed with a 1 mL micropipette. Another 1 mL of breaking buffer was added to the particles and removed as described above for multiple cycles until absolutely no white film was visible on the top of the supernatant. 400 μL of breaking buffer was then added, and the sample was transferred to a new 1.5 mL tube. Once again 1 mL of breaking buffer was added, vortexed, and centrifuged at 21,000 x g for 1 minute. The supernatant was removed, and the aptamer particles were resuspended in 1 mL of wash buffer.

ssDNA generation:

The aptamer particles were resuspended in 800 μL of 100 mM NaOH and incubated for 10 min at RT on a rotator. The aptamer particles were washed twice with 800 μL of 100 mM NaOH and then three times with 1 mL of wash buffer, briefly mixing between each wash. Finally, the aptamer particles were resuspended in 200 μL of wash buffer.

Aptamer particle quality control:

To ensure the successful synthesis and monoclonality of the aptamer particles, 1 μL of the aptamer particle solution, 98 μL of wash buffer and 1 μL of 100 μM Alex Fluor 647-modified RP was added to a 1.5 mL tube. The tube was incubated for 10 minutes at RT on a rotator and then washed once with 500 μL of wash buffer. The sample was then resuspended in 200 μL of wash buffer and run on a flow cytometer (BD Accuri C6 Plus). Monoclonality was assessed as previously described by our group^1^.

Protein labeling:

RA and RB were labeled using commercially available Dylight protein labeling kits (ThermoFisher Scientific). After the labeling reaction was complete, the samples were concentrated using a 3K Amicon Ultra-0.5 mL Centrifugal Filter (EMD Millipore) and then dialyzed in PBS using a 3.5K Slide-A-Lyzer MINI Dialysis Device (Thermo Fisher Scientific) overnight. The samples were then dialyzed in selection buffer for 4 hours and concentrated again using 3K Amicon filters. Finally, the absorbance of the DNA and the fluorophore were determined using a Nanodrop spectrophotometer, and the degree of protein labeling was determined using the formulas provided by the kit manufacturer.

Fetuin and asialofetuin were labeled by standard amine-NHS ester coupling. 0.3 mg fetuin and 0.3 mg of asialofetuin were separately resuspended in 0.05 M sodium borate buffer and incubated with 25 μg DyLight 650 N-NHS) ester or Dylight 532 NHS ester (Thermo Fisher Scientific) for 1 hour at RT. Labeled proteins were purified by dialysis (Slide A Lyzer MINI 10K, Thermo Fisher Scientific) overnight, and the degree of protein labeling was determined using a NanoDrop spectrophotometer.

MPPD screening protocol for RB-specific aptamers:

During each round of MPPD, ~10^8^ aptamer particles (the entire aptamer particle stock except for material used for quality control) was incubated with equimolar concentrations of labeled RA and RB (5 μM for the first round, 500 nM for the second round). Prior to incubation, the aptamer particles were resuspended in 500 μL of selection buffer and incubated for 15 minutes at RT on a rotator. After incubation, the beads were sonicated and resuspended with 250 μL of fluorescently labeled protein in selection buffer. The aptamer particles were incubated for 1 hour in the dark at RT on a rotator, washed once with 500 μL of cold selection buffer, resuspended in 3 mL of cold selection buffer, and put on ice. The aptamer particles were sorted on a BD Aria II FACS instrument using a 75-micron nozzle, and gating on the forward and side scatter was used to ensure that only singlet beads were analyzed. Around 0.1–0.3% of the singlet aptamer population was collected that displayed the greatest shift in RB binding (FITC channel) without any increase in RA binding (APC channel). The aptamer particles were kept at 4 °C in the FACS sample chamber. After sorting, the collected aptamer particles were transformed back to natural dsDNA as previously described^1^.

Finally, each aptamer pool was assessed for enrichment of RA and RB binders. 1 μL of the aptamer particle solution was added to fluorescently-tagged RA and/or RB in selection buffer in a total reaction volume of 25 μL. The samples were incubated in the dark for at least 45 minutes at RT on a rotator. Each sample was individually washed with 500 μL of cold selection buffer, resuspended in 200 μL of cold selection buffer and run on a benchtop cytometer (BD Accuri C6 Plus) at a slow flow-rate. The singlet population was identified by gating in the side-scatter and forwards-scatter plots, and the fluorescent signal of the population was then analyzed to assess binding.

MPPD screening protocol for fetuin-specific aptamers

FP-modified beads were synthesized as described above. Aptamer particles were generated by emulsion PCR using an oil phase composition of 4.5% Span 80, 0.45% Tween 80, and 0.05% Triton X-100 in mineral oil. The aqueous phase consisted of 1x KOD XL DNA polymerase buffer, 50 U KOD XL DNA polymerase, 0.2 mM dATP, 0.2 mM dGTP, 0.2 mM dCTP, 0.2 mM C8 alkyne dUTP, 10 nM Fet FP, 1 μM Fet RP, 1 pM template DNA, and 12 μl FP beads. 1 mL of aqueous phase was added to 7 mL of oil phase and emulsified at 620 rpm for 5 min in an IKA DT-20 tube using the IKA Ultra-Turrax device. The emulsion was pipetted into 100 μL reactions in a 96-well plate and amplified using the protocol: 95 °C for 3 min, 39 cycles of 96 °C for 15 s, 57 °C for 30 s, 74 °C for 1 min and final step at 72°C for 5 min. Emulsion cleanup and aptamer particle quality control cleanup were performed as described for RNAse A/B. Aptamer particles were transformed into non-natural aptamer particles using CuAAC click chemistry as follows. Aptamer particles recovered from the emulsion PCR were washed twice with 200 μL 1X PBS and resuspended in a 200 μL solution containing 1X PBS, 5 μL of a pre-prepared mixture of 0.1 M CuSO4/0.2 M tris(3-hydroxypropyltriazolylmethyl)amine (THPTA), and 10 mM tryptophan azide, and H_2_O to the final volume. A 50 mM solution of sodium ascorbate was freshly prepared in H_2_O, and 20 μL was added to the reaction mixture for a final concentration of 5 mM. The solution was degassed using N_2_ for 5 minutes, then reacted for 40 min with rotation at room temperature. ssDNA generation was accomplished by resuspending the particles in 200 μL 0.1 M NaOH and incubating for 10 min at RT. Beads were washed three times and resuspended in fetuin selection buffer.

One round of MPPD was performed with Dylight 650-labeled fetuin and Alexa Fluor 532-labeled asialofetuin. Prior to sorting, a flow cytometry binding assay was performed with multiple concentrations of fetuin (10, 50, 100, 200 nM) to determine which resulted in sufficient binding to the target. A second binding assay was then performed using the concentration of fetuin determined in the first assay in the presence of multiple concentrations of asialofetuin (10, 50, 100, 200 nM). For the assay, 1 μL of the aptamer particle solution was added to a solution containing the labeled protein targets at the specified final concentrations in a total reaction volume of 50 μL of 1X fetuin selection buffer and incubated in the dark for at 1 hr at RT on a rotator. Samples were then washed with 200 μL and resuspended in cold 1X fetuin selection buffer for analysis. The optimal concentration of asialofetuin was determined as the highest concentration at which >1% binding to Dylight 650-fetuin remained. For sorting, aptamer particles were folded in 1 mL fetuin selection buffer and then incubated with 100 nM Dylight650-labeled fetuin and 200 nM Alexa Fluor 532-labeled asialofetuin on a rotator in the dark for 1 h at room temperature. The beads were washed twice and resuspended in 1 mL cold 1X selection buffer and then analyzed using a BD FACS Aria III. The sort gate was set to collect 0.3% of aptamer particles that showed high specificity for fetuin by identifying those particles with the greatest shift in the APC channel (fetuin) and no overlap in the PE channel (asialofetuin). After sorting, the collected aptamer particles were resuspended in 20 μL PBS and the aptamers were amplified by PCR using the conditions above.

High-throughput sequencing protocol:

For each aptamer pool sequenced, adaptor primers were first added. 10 ng of dsDNA was subjected to eight cycles of PCR. A 2x GoTaq Master Mix was used (Promega, M7132) with 1 μM of each primer (100 uL reaction volume). The sequencing primers were added by using a Nextera XT kit (Illumina) and following the provided instructions. Samples were quantified using a Qubit fluorometer and sent to the Stanford Functional Genomics Facility (SFGF) for sequencing on an Illumina MiSeq.

Family analysis of sequencing data:

All analysis was conducted in Python. First, the number of replicates were counted for each sequence, and sequences with N or more replicates were selected (N = 3 for RB selection, and n = 2 for fetuin). Using the levenshtein_distance function from the Levenshtein Python library, the edit distance between all the sequences was calculated and stored. The sequences were then clustered into families of edit distance of 3 or less, and sorted based on the abundance of family members.

Synthesis of aptamer particles:

Aptamers were coated onto beads by preparing a 100 μL PCR reaction consisting of 10 μL 10X KOD XL buffer, 1 μL dNTP mix of 10 mM of each nucleotide (containing the modified-nucleotide analog), 0.5 μL of 10 μM FP, 2.5 μL of 100 μM RP, 5 μL of 10 nM aptamer template, 8 μL of FP beads, 71 μL of water, and 2 μL of KOD XL polymerase. 30 PCR cycles were conducted. The beads were washed and converted to ssDNA as described above for the ePCR protocol, and resuspended in 50 μL of selection buffer prior to storage at 4 °C.

Affinity measurements:

A 10 μL binding reaction was setup with 2 μL of the aptamer particle solution, the required volume of the fluorescently-labeled protein stock in selection buffer, and then brought up to 10 uL in selection buffer. For the RB competition assays, the competitor molecule (concentrated stock solution in selection buffer) was added to the reaction mixture immediately before adding the fluorescently-labeled protein stock. The samples were incubated on a rotator at RT for 45 minutes and then put on ice. 2 μL of the reaction was added to 200 μL of cold selection buffer and put on a magnet for 1 minute. The supernatant was discarded, and the particles were washed with 500 μL of cold selection buffer and then resuspended in 200 μL of cold selection buffer. The sample was gently mixed via pipette and then immediately run on a flow cytometer (BD Accuri C6 Plus).

PNGase F deglycosylation of RNase B:

5 μL of Remove-iT PNGase F (NEB), 3 μL of selection buffer, and 12 μL of a 40 μM RB stock in selection buffer were added to a 1.5 mL tube and incubated at 37 °C for 15 hours while shaking at 350 RPM. Magnetic chitin beads (NEB) were then used to remove the PNGase F, which contains a chitin binding domain (CBD). 50 μL of chitin beads were washed twice with 200 μL of selection buffer, and then combined with the deglycosylation reaction. The mixture was incubated for 30 minutes on a rotator at RT, and then the supernatant was collected. The deglycosylated RB was stored at 4 °C.

**SI Tables:**

**Figure 2:** Overview of particle display selection scheme. 1. Magnetic beads are coated with forward primers 2. Emulsion PCR creates aptamer particles that each display many copies of a single aptamer sequence. The modified base 5-Indolyl-AA-dUTP is introduced along with dATP, dCTP, dGTP. 3. Emulsions are broken and cleaned up 4. Aptamers are made single stranded 5. Aptamer particles are incubated with RNase B and RNase A. Each protein is labeled with an orthogonal fluorophore. 7. FACS screening separates aptamer particles that bind with high specificity to RNase B. This is done by collecting events in quadrant IV of the FACS plot. 8. Natural DNA is amplified from the enriched aptamer particles. 9. After several rounds of Click-MPPD, aptamer sequences are identified through high throughput sequencing.

| **Name** | **Sequence** |
| --- | --- |
| i-1 | ZZZGZGCGCZZCAGZCAAZZGGCAZZCGZGAAZAZCZCCZZZZACGACAC |
| i-2 | ZCGGZGGACGCCZCZZGAZCGAGZZZZGGZCAZGZGZZZACGZAZGZZZC |
| i-3 | GCGGZZCZZACZCAGCCCZAZGCZACGACACZCZZZCZZACCCCAACCZA |
| i-4 | ZZZGZGCGCZZCAGZCAAZZGGCAZZCGZGAAZAZCZCCZZZZZACGACA |
| i-5 | CGGZZCZZACZCAGCCCZAZGCZACGACACZCZZZCZZACCCCAACCZAC |
| i-6 | AZCZCGZGZGZZZGGZZZZAGAZZZZZGGAAAZZAZGAZZCCZGZZGGZC |
| i-7 | ZAGCAZGZCAGZCAACCZZCAGZAZGCZCGZZGZGAZZZZZZZCGCAZAC |

**Table S1:** RB aptamer sequences selected from HTS data. Primer regions not shown. The indole-modified dUTP analog is represented by the letter Z.

| **Name** | **Sequence** |
| --- | --- |
| f-1 | CAAAGCCACZGAACCACCCGGCGGAAAAAGAGAZGGAGCA |
| f-2 | AGAAAAGCGAGAAACCGGAGGGAGCGCGCGGCAACCGACA |
| f-3 | CAAAAAGAAACCAGCAAZCAAAACCCGCAAGCAGCAAZAG |
| f-4 | CGCCAAAGCZCGAZCAAZAACCGAAAGAAAAGGCACAZCA |
| f-5 | CCGGGCCACGAACZCAAZGGAGACGCACGCCAAACACGGA |
| f-6 | CAGAAAAAGGGAGZGGGZGGGZCGGGCAGAGGAAGGGGAG |
| f-7 | CAGAAGAAGZGAGZGGGZGGGZCGGGCAGAGGAAGGGGAG |

**Table S2**: Fetuin aptamer sequences selected from HTS. Indole-modified dUTP analogs are represented by the letter Z.

**SI Figures:**

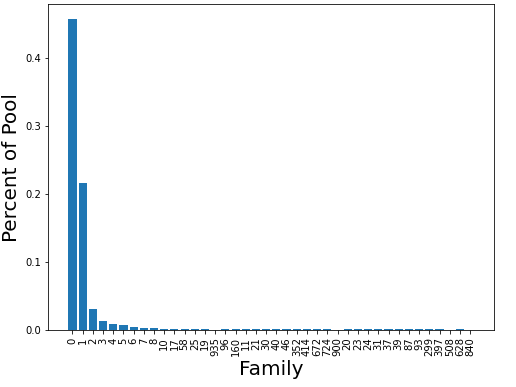

**Figure S1:** Abundance analysis of RB aptamer families. Aptamer families were identified from the HTS data after MPPD selection and were defined as sequences with a Levenshtein distance < 3 as described in the methods section.

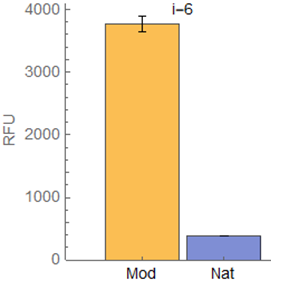

**Figure S2**: Binding of base-modified and natural DNA versions of aptamer i-6 to 100 μM fluorescently-labeled RB. Experiments were conducted in triplicate and the error bars represent the standard deviation of the median RFU values.

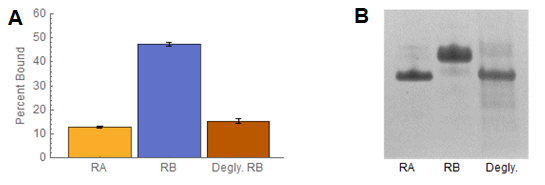

**Figure S3: A)** Bead-based binding assays with 20 μM fluorescently labeled RA, RB, and deglycosylated RB. Experiments were conducted in triplicate and median RFU values were used. Error bars represent a single standard deviation. **B)** SDS-PAGE analysis of RA, RB, and RB after enzymatic cleavage of the glycan.

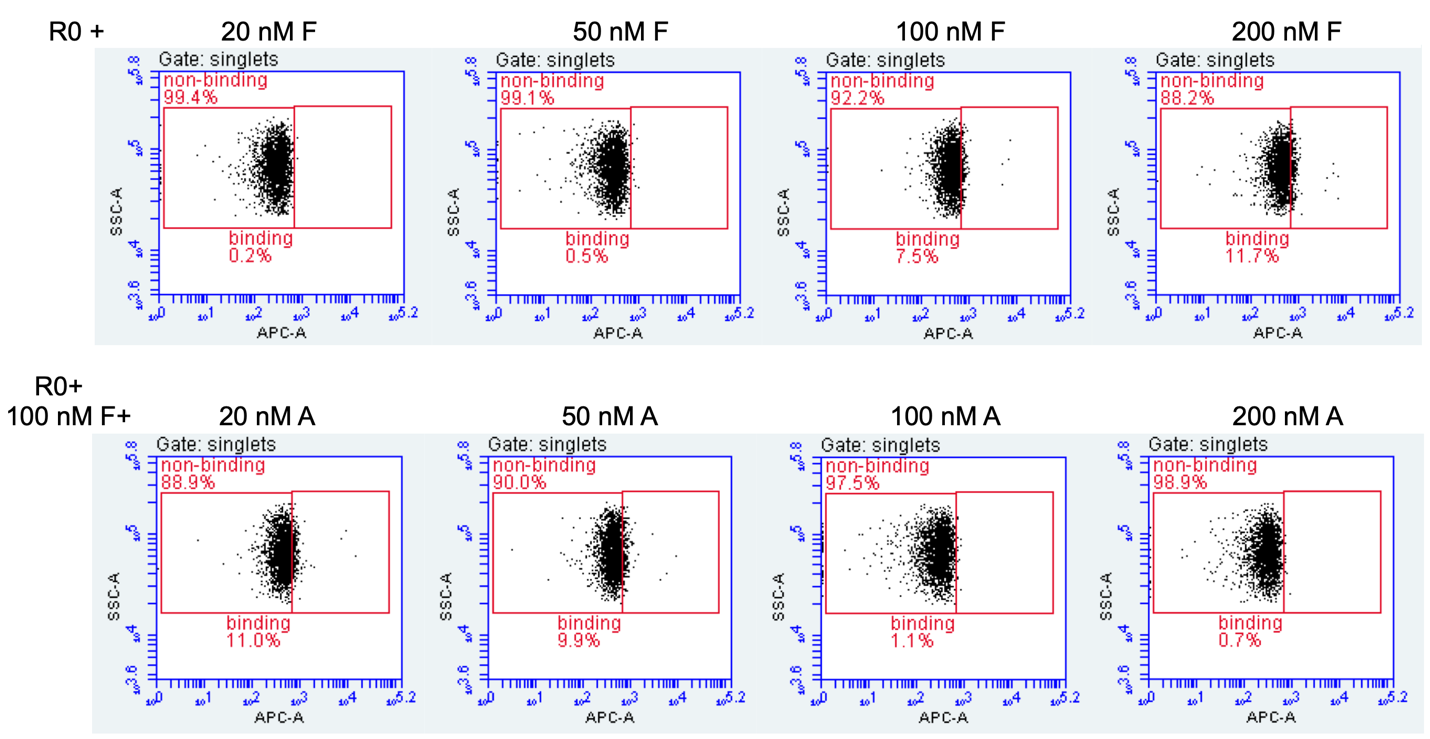

**Figure S4: Flow cytometry based binding assay of R0 (pre-enriched) library prior to PD screening.** An initial assay was performed to evaluate binding of the library to fetuin alone to determine which concentration resulted in a sufficient shift in the APC channel. A concentration of fetuin was chosen from this (100 nM) and a second assay was performed to determine how much binding to Dylight-650 labeled fetuin was retained in the presence of increasing Alexa Fluor 532-labeled asialofetuin. Final concentration of 100 nM fetuin and 200 nM asialofetuin were chosen for two-color based FACS.
